# Interhemispheric connections between olfactory bulbs improve odor detection

**DOI:** 10.1101/767210

**Authors:** Florence Kermen, Emre Yaksi

**Author notes:** Corresponding authors &.

## Abstract

Interhemispheric connections enable interaction and integration of sensory information in bilaterian nervous systems and are thought to optimize sensory computations. However, the cellular and spatial organization of interhemispheric networks as well as the computational properties they mediate in vertebrates are still poorly understood. Thus, it remains unclear to which extent the connectivity between left and right brain hemispheres participates in sensory processing. Here, we show that the zebrafish olfactory bulbs (OBs) receive direct interhemispheric projections from their contralateral counterparts in addition to top-down inputs from the contralateral zebrafish homolog of olfactory cortex. The direct interhemispheric projections between the OBs reach peripheral layers of the contralateral OB and retain a fine-grained topographic organization, which directly connects similarly tuned olfactory glomeruli across hemispheres. In contrast, interhemispheric top-down inputs consist of diffuse projections that broadly innervate the inhibitory granule cell layer. Jointly, these interhemispheric connections elicit a balance of topographically organized excitation and non-topographic inhibition on the contralateral OB and modulate odor responses. We show that the interhemispheric connections in the olfactory system enable the modulation of odor response and improve the detection of a reproductive pheromone, when presented together with competing complex olfactory cues, by boosting the response of the pheromone selective neurons. Taken together, our data shows a previously unknown function for an interhemispheric connection between chemosensory maps of the olfactory system.

## INTRODUCTION

In bilaterians, information detected by parallel sensory channels is topographically represented onto sensory regions located in mirrored positions of the brain hemispheres. Communication between bilateral sensory regions with similar functions is mediated by a dense network of fibers projecting contralaterally between brain hemispheres, which are organized into commissural tracts [1–4]. These interhemispheric connections are present in invertebrates [5,6] and vertebrates [7,8]. Different theories exist, pertaining to how they contribute to neural computations, in particular with regards to their inhibitory or excitatory effect [9,10]. Interhemispheric inhibition has been suggested to facilitate visual [11,12] and somatosensory processing [13,14] by increasing the perceptual threshold in the hemisphere contralateral to the stimulated hemisphere, and thereby shifting attention to the relevant input. A functional diversity of interhemispheric connections is present in the auditory system, where both inhibitory and excitatory interhemispheric interactions have been observed [15,16]. The synchronization between auditory hemispheres might support figure/ground segregation, whereas interhemispheric inhibition amplifies binaural activation delay and could underlie stereo-hearing [17]. On the other hand, information sharing between hemispheres through excitation in the primary visual cortex enables binocular vision supporting depth perception [18,19]. It also avoids the dissociation between lateralized associative centers, by transferring unilaterally learned information to the opposing hemisphere. For example, the restricted lesion of fibers linking the somatosensory cortices blocks the transfer of texture learning from one paw to the other in monkeys [20]; and disconnecting the fibers responsible for the visual interhemispheric crosstalk prevents the transfer of a learned visual discrimination task from one hemifield to the other [21]. Despite extensive studies, the precise cellular-level mapping of interhemispheric projections linking bilateral sensory regions, as well as the functional properties they mediate remains largely elusive.

In the olfactory system, the activation of receptors located in spatially segregated bilateral nostrils is transformed through ipsilateral projections into topographically organized, mirror-symmetric sensory maps in the OBs [22–24]. However, these segregated olfactory processing channels were also shown to exhibit prominent interhemispheric interactions in several taxa. For example in fruitflies, sensory neurons project bilaterally to the antennal lobes [25], and were suggested to improve the signal quality by pooling multiple input channels from both antennae [26]. Rodents are able to locate an odor source using bilateral sampling [27,28], which depends on intact interhemispheric communication [27]. A multisynaptic system links the left and right OBs via the anterior olfactory nucleus (AON) [29], a circuit which was proposed to support interhemispheric transfer of olfactory memory [30]. Beyond such connections through higher olfactory areas of vertebrates, direct projections were shown to link both OBs in teleost fish [31]. It is however less clear, how all these interhemispheric connections between olfactory processing channels across the vertebrate brain work in parallel, and contribute to olfactory computations. The function of interhemispheric olfactory connections is perhaps best conceptualized when asking what could be the advantage for vertebrates to maintain redundant olfactory maps in each hemisphere. Indeed, unilateral olfactory input is sufficient to detect, identify and spatially locate odors [32,33]. Yet, bilateral sensory maps linked by interhemispheric connections could confer additional properties akin to visual depth perception [34] and fine sound source localization [35]. In line with this, prior evidence suggest that bilateral olfactory inputs improve the speed and the accuracy of odor tracking in a variety of animal species [28,32,33,36]. The neural circuits serving the improved chemotaxis are currently unknown, however mechanisms have been proposed, including increased signal-to-noise ratio [32] or interhemispheric inhibition [37]. Alternatively, redundant information could prove essential when a sensory organ is (partially) obstructed, as can be the case in both rodents and humans [38,39]. Interhemispheric transfer of unilaterally experienced odorant information through AON would then help maintain both olfactory maps in the cortex up to date, even when one of the sensory organs is offline [30,39,40]. Despite these studies, the precise neural circuit that links the two olfactory hemispheres and how this circuit contributes to olfactory computations is yet to be further elucidated.

To address this, we used an ex vivo preparation of the adult zebrafish brain and characterized interhemispheric connections between the primary brain regions processing odor information, the OBs. First, we showed that the olfactory glomeruli receive direct and spatially organized mitral cell projections from similarly tuned glomeruli at the contralateral olfactory bulb. Additionally, we showed that the OB interneurons receives top-down projections from the contralateral telencephalic olfactory centers. Second, we investigated the function of these interhemispheric connections by using two-photon calcium imaging in the OB. We found that interhemispheric connections elicit a balance of topographically organized odor-specific excitation and non-topographically organized inhibition on the contralateral OB. Our results revealed that strong contralaterally evoked-activity can modulate odor responses in an intensity dependent manner. Finally, we demonstrated that the excitatory and inhibitory interhemispheric interactions jointly improve the detection of a reproductive pheromone within a background of olfactory noise elicited by food odors, by specifically boosting the response of the pheromone responding OB neurons. Taken together, our findings show a previously undescribed function for the interhemispheric olfactory connections for improving the detection of sensory stimuli within a noisy background.

## RESULTS

### The olfactory bulbs are connected through direct and indirect interhemispheric projections

In vertebrates, olfactory information from one nostril is sent to the ipsilateral OB. It is subsequently passed on to higher olfactory areas primarily on the ipsilateral side[41]. In the current study, we used an ex vivo explant preparation of the adult zebrafish brain [22,42,43] to investigate how bilateral olfactory bulbs are interconnected across the brain hemispheres. We first investigated whether mitral cells directly project to the contralateral OB (interbulbar projections). To do so, OB neurons were labeled by electroporation of tetramethylrhodamine-dextran. Two-photon microscopy was then used to image the labeling sites as well as the axonal projections to the contralateral OB (Figure 1A-B). Mitral cells axons exited the ipsilateral OB through the medial and lateral olfactory tracts and mostly terminated in the ipsilateral Dp (dorsal part of the dorso-lateral pallium), which is a zebrafish telencephalic area homologous to the rodent primary olfactory cortex [44] (Figure 1A). Interestingly, we also observed that axons coursing through the medial olfactory tract crossed to the contralateral hemisphere at the level of the anterior commissure (Figure 1A). These axons terminated at the peripheral layers of the contralateral olfactory bulb, where the glomeruli and the mitral cells are located (Figure 1A-C).

**Figure 1:**
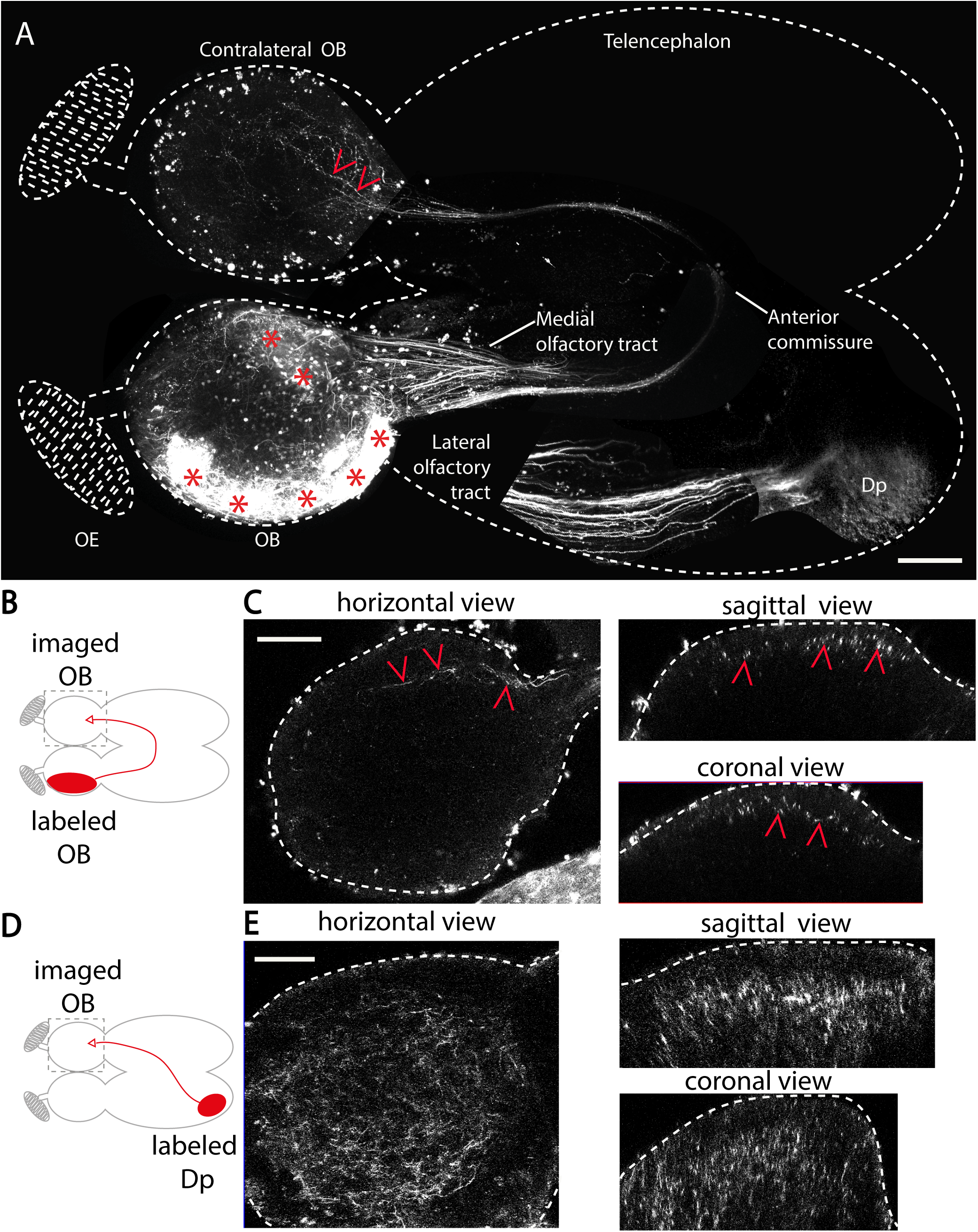
Zebrafish olfactory bulbs are connected by two interhemispheric pathways. (A). Two-photon microscopy images showing unilateral dye labeling of olfactory bulb cells in an adult zebrafish forebrain. This image is reconstructed by juxtaposing several partially overlapping images. Labeled mitral cells axons cross the midline at the level of the anterior commissure and terminate in the contralateral olfactory bulb. Dense ipsilateral axonal projections to Dp, the fish homolog of the olfactory cortex, course through the lateral olfactory tract. Red asterisks indicate dye electroporation sites in the olfactory bulb and red arrowheads point at axonal projections in the contralateral olfactory bulb. (B). Schematic of dye labeling in the olfactory bulb. The filled red ellipse indicates the labeling site. The grey dashed square indicates the imaging zone in C. (C). Horizontal, sagittal and coronal views of mitral cell axons projecting to the contralateral olfactory bulb in a representative brain (n=12). (D). Schematic of dye labeling of Dp. The grey dashed square indicates the imaging zone in E. (E). Horizontal, sagittal and coronal views of Dp projections to the contralateral OB in a representative brain (n=5). Scale bars represent 100 µm. Dp: dorsal part of the dorso-lateral pallium; OE: olfactory epithelium; OB: olfactory bulb.

The vertebrate OB is innervated by centrifugal fibers originating from the ipsilateral telencephalon, where higher olfactory centers are located [45–48]. To investigate the extent of contralateral inputs received by the OB from higher olfactory centers, we asked whether top-down projections from Dp reached the contralateral OB in adult zebrafish. To do so, we unilaterally labeled Dp neurons using local dye electroporation (Figure 1D). Neural processes extending from the Dp crossed the anterior commissure and diffusely innervated the core layer of the contralateral OB (depth> 50µm, Figure 1E), where the granule inhibitory interneurons are located. Altogether, our results revealed that the zebrafish OBs are connected through at least two major types of interhemispheric connections. A direct and topographically organized mitral cell projection terminates on the peripheral layer of the contralateral OB. An indirect connection through the zebrafish homolog of the olfactory cortex, Dp, diffusely terminates on the granule cell layer of the contralateral OB.

### The direct interbulbar projections retain a fine scale chemotopic organization

In the zebrafish OB, odors with similar chemical properties are encoded by the activation of spatially nearby domains forming a chemotopic map [22] that is conserved across individual animals [49,50]. For example the neurons responding to amino acids, which are feeding cues, are located in the lateral OB [51], while neurons responding to bile acids, which are putative social cues, are located in the medial OB [22,51]. We thus asked whether projections of mitral cells at the contralateral OB are topographically organized. Labeling mitral cells located in the medial OB revealed projections terminating at the medial domain of the contralateral OB (Figure 2A-B), thus mirroring their cell body location. Similarly, labeling mitral cells at the lateral OB revealed projections terminating at the lateral domain of the contralateral OB (Figure 2C-D). These results suggested that direct interbulbar projections are topographically organized with respect to the functional domain they originate from.

**Figure 2:**
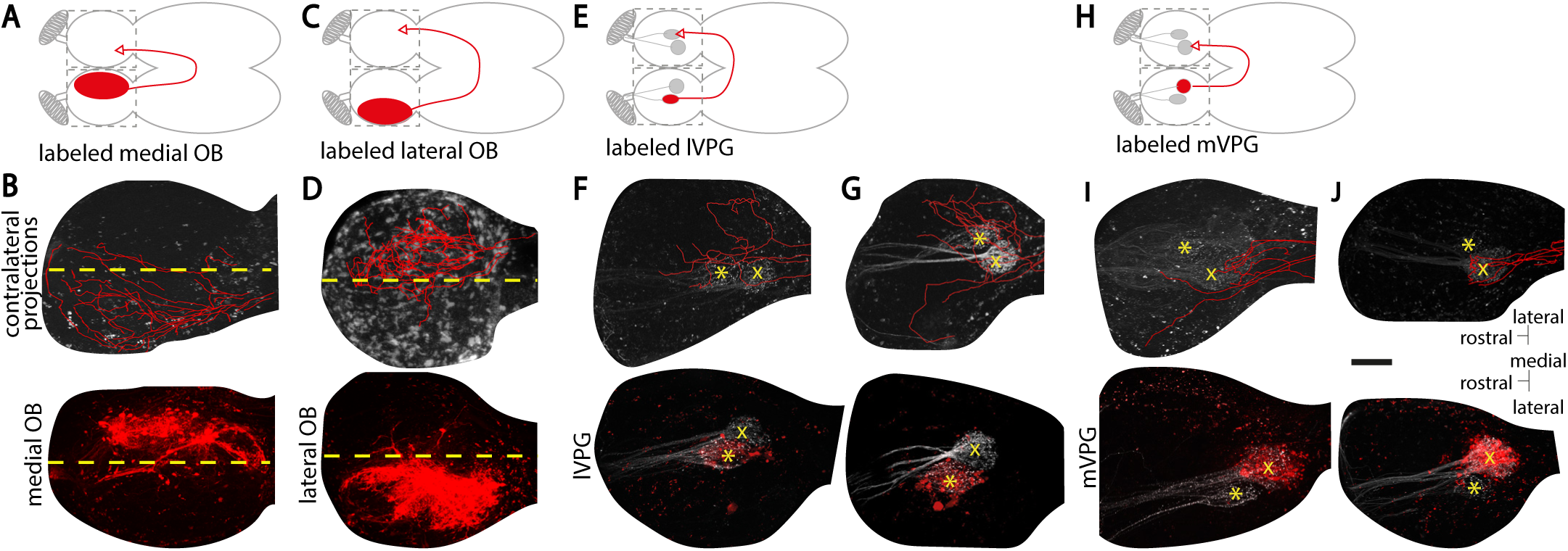
The direct interbulbar connections are topographically organized and glomeruli specific. (A). Schematic of the dye labeling in the medial olfactory bulb. The filled red ellipse indicates the labeling site. The grey dotted squares indicate the two imaging fields. (B). Two-photon microscopy images showing the labeled neurons in the medial olfactory bulb (bottom) and their reconstructed axonal projections (red) to the contralateral medial olfactory bulb (top, n=1). The yellow dotted line separates each olfactory bulb into a medial and a lateral half. (C). Schematic of the dye labeling in the lateral olfactory bulb. (D). Two-photon microscopy images showing the labeled neurons in the lateral olfactory bulb (bottom) and their reconstructed axonal projections (red) to the contralateral lateral olfactory bulb (top) in a representative brain (n=7). (E). Schematic of the dye labeling of the lVPG. (F & G). Two-photon microscopy images showing the labeled neurons innervating the lVPG (bottom) and their reconstructed axonal projections (red) to the contralateral olfactory bulb (top, n= 2). (H) Schematic of the dye labeling of the mVPG. (I & J). Two-photon microscopy images showing the labeled neurons innervating the mVPG (bottom) and their reconstructed axonal projections (red) to the contralateral olfactory bulb (top, n= 2). The lVPG is indicated by a yellow asterisk and the mVPG by a yellow cross. Scale bar represents 100µm. lVpG: lateral ventro-posterior glomerulus; mVpG: medial ventro-posterior glomerulus; OB: olfactory bulb.

Next, we tested whether the direct interbulbar projections can specifically link mirror-symmetric glomeruli with similar odor tuning (iso-functional glomeruli) across hemispheres. To do this, we labeled mitral cells innervating a genetically identified glomerulus lVPG (lateral ventro-posterior glomerulus, Figure 2E), which can be visualized in similar locations in both OBs (Koide et al., 2009). Examination of the electroporation site confirmed that the labeled neurons indeed extended dendrites connecting to the lVPG glomerulus (Figure 2F-G, bottom panels), but spared the adjacent and medially positioned mVPG (medial ventro-posterior glomerulus) thus emphasizing the spatial specificity of the electroporation [52]. We observed that lVPG mitral cells sent interhemispheric projections that terminated next to the contralateral lVPG, as well as in nearby glomeruli (Figure 2F-G, top panels). We repeated this experiment by labeling the mitral cells of the mVPG (Figure 2H). We observed that the axons of mVPG mitral cells projected to the mVPG at the contralateral olfactory bulb (Figure 2I-J). Our results showed that direct interbulbar projections across the brain hemispheres are topographically organized and glomeruli specific.

### Interhemispheric connections between olfactory bulbs modulate odor responses

The presence of these extensive interhemispheric connections raised the possibility that olfactory information in one hemisphere can influence odor processing in the contralateral OB. To test this hypothesis, we measured odor-evoked responses using two-photon calcium imaging in zebrafish expressing GCaMP6s under the elavl3 promoter, primarily in mitral cells [42,53], while electrically stimulating the contralateral olfactory nerve at different intensities using a glass microelectrode (Figure 3A). As expected, food odor activated a large number of ipsilateral mitral cells (Figure 3C-E, left panels). Low intensity microelectrode stimulation of the contralateral olfactory nerve led to modulation of odor response in very few ipsilateral mitral cells (Figure 3C & F, right panels). However, as we increased the contralateral stimulation to medium and strong intensities, more ipsilateral mitral cells showed increased odor responses (Figure 3D,G,E,H). Importantly, to verify that contralateral stimulation did not directly activate ipsilateral neurons through volume transmission, we repeated the same experiment after sectioning contralateral mitral cells’ axons at the level of the olfactory tract (Figure 3B). In that case we did not observe modulation of mitral cell responses, even when using the strongest microstimulation intensity (Figure 30-R), indicating the necessity of intact contralateral olfactory inputs. We next calculated the contribution of contralateral inputs to odor responses by measuring the difference of amplitude between the ipsilateral odor responses with and without the contralateral microelectrode stimulation (Figure 3I-K). We found significant and mainly excitatory modulation of odor responses by medium and strong contralateral inputs (Figure 3M-N). Our results showed that interhemispheric connections across the OBs can dynamically modulate the odor response in the contralateral olfactory bulb.

**Figure 3:**
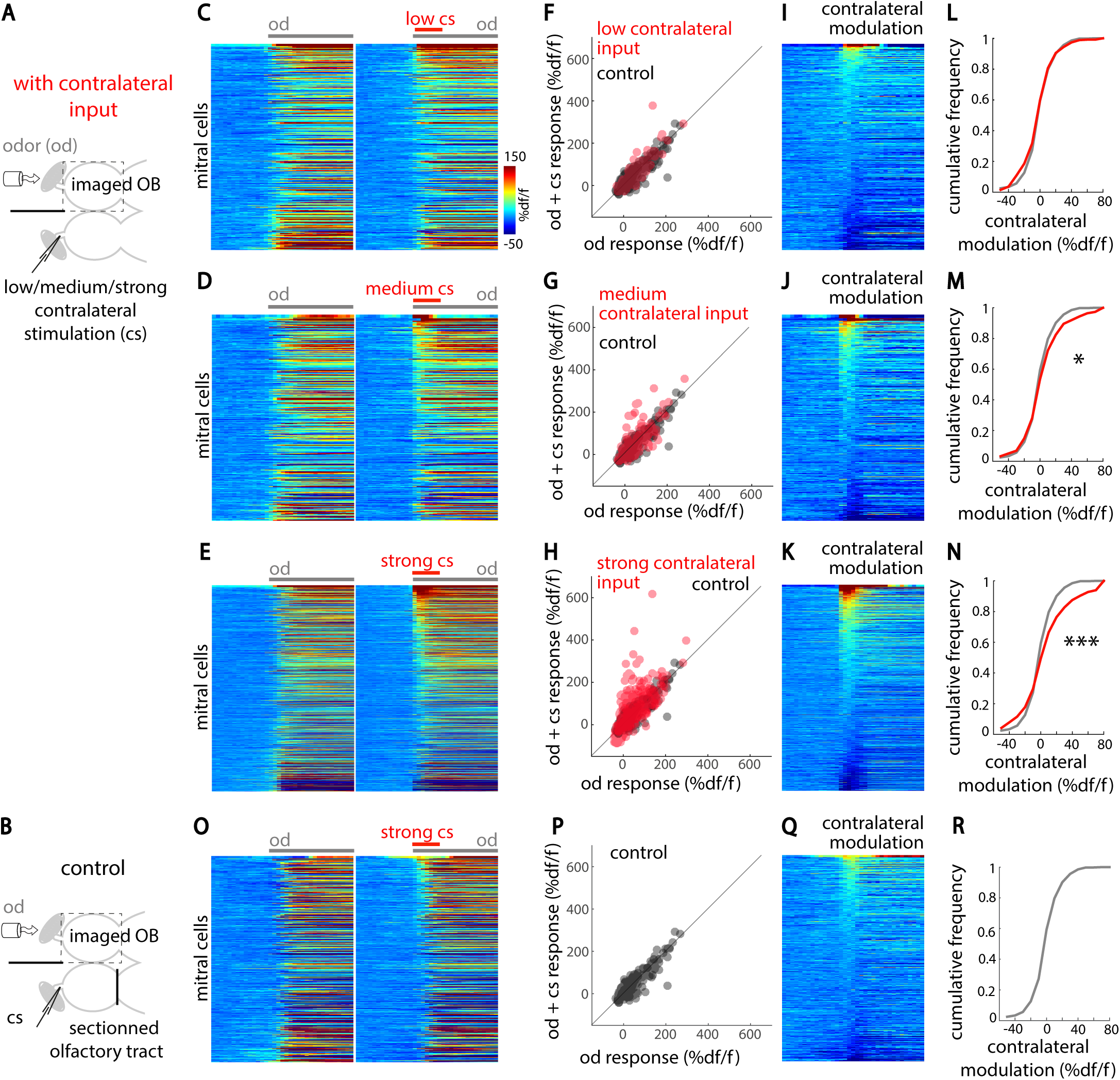
Interhemispheric connections modulate odor-evoked responses in the contralateral olfactory bulb. (A). Illustration of the experimental setup depicting a adult zebrafish brain explant, expressing GCaMP6s in mitral cells. A tube delivers food odor (od) selectively to the ipsilateral nostril. Contralateral stimulation (cs) of varying intensities (low, medium and strong, see Methods) are delivered using a glass microstimulation electrode inserted in the contralateral olfactory nerve. (B). The contralateral olfactory tract, which enables interhemispheric olfactory connections, is sectioned in control conditions. (C-E & O). Mean odor response time course of all mitral cells recorded in the ipsilateral olfactory bulb following odor presentation (od, grey bar) or odor combined with simultaneous contralateral microelectrode stimulation (od + cs; cs indicated by red bar) for low, medium and strong stimulation intensities and the control condition, respectively. For each condition, mitral cells from multiple animals are pooled and sorted by the amplitude of contralateral modulation ([od + cs] - od). (F-H & P). Average mitral cells responses to odor (red) are plotted against their response to odor combined with contralateral microstimulation for low, medium and strong contralateral microstimulations (in red) and the control condition (in black), respectively. Mitral cells above the unity line are positively modulated by contralateral inputs. (I-K & Q). The contralateral modulation ([od + cs] - od) is displayed for all mitral cells for low, medium and strong contralateral microstimulations and the control condition, respectively. (L-N & R). The cumulative frequency distribution of the contralateral modulation is displayed for all mitral cells for low, medium and strong contralateral microstimulations (in red) and the control condition (in black), respectively. The number of mitral cells were as follow: low cs: 335 cells in 3 fish; medium cs: 335 cells in 3 fish; strong cs: 850 cells in 6 fish; control cs: 477 cells in 2 fish; non-significant; * p<0.05; *** p<0.001; two-sample two-tailed Kolmogorov-Smirnov test. cs: contralateral microstimulation; od: food odor extract; OB: olfactory bulb.

### Natural odorselicit odor-specific and chemotopically organized excitation in the contralateral olfactory bulb

Having shown that contralateral olfactory nerve electric stimulation can modulate odor responses, we further investigated whether odor stimulation would also elicit activity in the contralateral OB. To assess this, we sectioned the olfactory nerve of the contralateral OB and delivered odors to the ipsilateral side with an intact olfactory nerve. By performing two-photon calcium imaging, we compared the odor-evoked mitral cell activity in the ipsilateral OB (Figure 4A) with the activity in the contralateral deafferented OB (Figure 4B) in response to amino acids, bile acids and a reproductive fish pheromone, prostaglandin 2α (pgf2α). We observed neural responses to all odors, both in the afferented (Figure 4C-D) and in the deafferented OB neurons (Figure 4E-F), which showed that natural odors indeed elicit excitatory responses in the contralateral OB. As expected, there were more ipsilateral than contralateral odor-responsive mitral cells (Figure 4G). Importantly, no odor-induced activity was seen in bilaterally deafferented OBs (control condition, Supplementary Figure 1). The results showed that natural odors can elicit activity not only in the ipsilateral but also in the contralateral OB.

**Figure 4:**
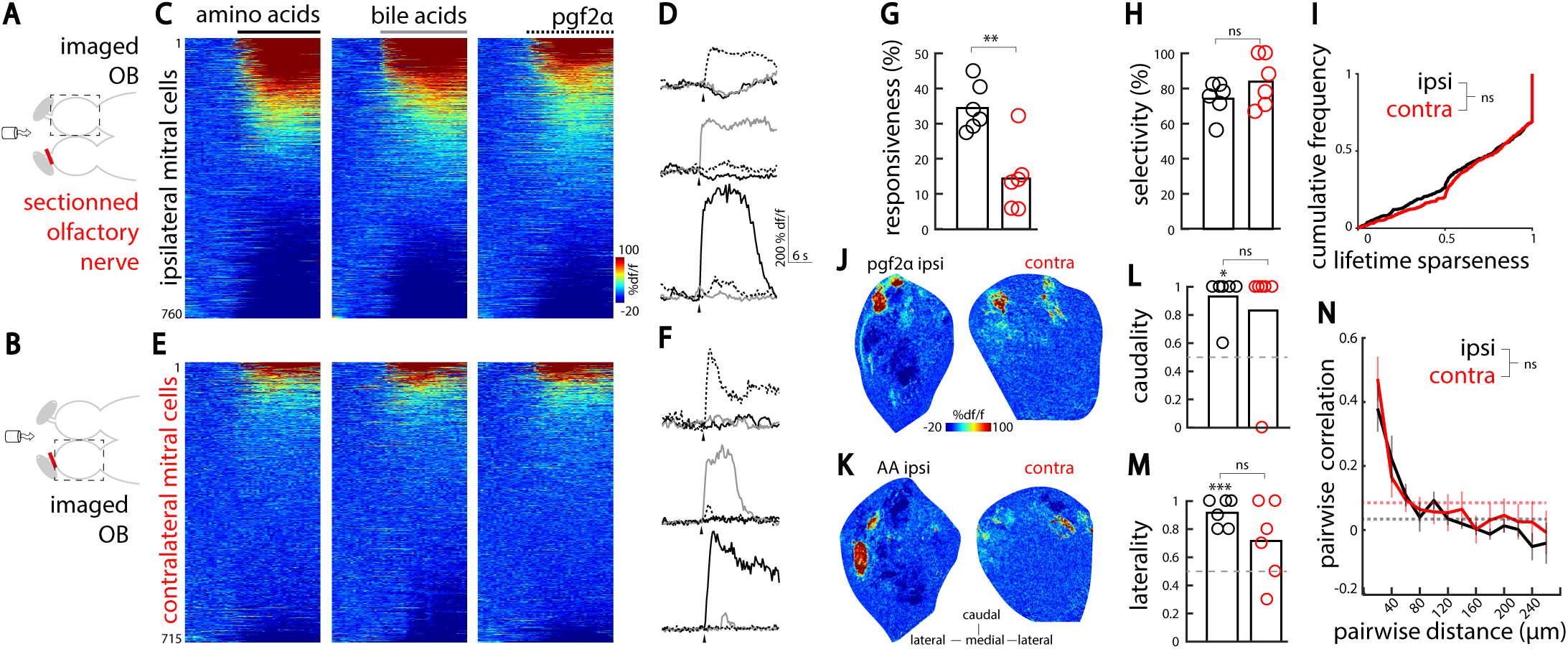
Natural odors elicit odor-selective and chemotopically organized responses in the contralateral olfactory bulb. (A). Illustration of the experimental setup depicting an adult zebrafish brain explant, expressing GCaMP6s in mitral cells. A tube delivers the olfactory cues to the nostrils. The stimuli used were amino acids, bile acids and pgf2α. The olfactory nerve was sectioned (red mark) on contralateral olfactory bulb. Mitral cell odor responses are recorded in the olfactory bulb with intact olfactory nerve, using two photon calcium imaging. (B). Illustration of the experimental setup depicting an adult zebrafish brain explant. Mitral cell odor response are recorded in the deafferented OB, using two photon calcium imaging. (C). Odor response time course of all ipsilateral mitral cells pooled and sorted by their response amplitude (760 cells in 6 fish). (D). Odor response time course of three representative odor-selective ipsilateral mitral cells. Arrowheads indicates odor onset. (E). Odor response time course of all contralateral mitral cells pooled and sorted by decreasing responsiveness (715 cells in 6 fish). (F). Odor response time course of three representative odor-selective contralateral mitral cells. Arrowheads indicates odor onset. (G). The percentage of mitral cells responding to at least one olfactory cue is larger in the ipsilateral OB (black) than in the deafferented contralateral OB (red) (**p<0.01; Student’s t-test). (H). The percentage of mitral cells responding to only one of the presented odors was similar in the ipsilateral (black) and contralateral OB (red) (Student’s t-test). (I). The cumulative frequency of lifetime sparseness for all mitral cells was similar in both olfactory bulbs. two-sample two-tailed Kolmogorov-Smirnov test. (J). Ipsilateral (left) and contralateral (right) OBs responses to pgf2α in the same fish (representative planes located 50 µm deep from the ventral OBs surface). (K). Ipsilateral (left) and contralateral (right) OBs responses to amino-acid mix in the same animal than in J (representative planes located 90 µm deep from the ventral OBs surface). (L). Spatial distribution of mitral cells responding selectively to pgf2α, along the rostro-caudal axis in both OBs for all fish. Values close to 1 indicate caudal locations, whereas values close to 0 indicate rostral locations. Note the caudal location of pgf2α compared to the random distribution indicated by the gray dotted line (*p<0.05; Mann Whitney U test). Pgf2α-selective cells are similarly located in the contralateral and ipsilateral OBs (Mann Whitney U test). (M). Location of mitral cells responding selectively to amino-acids, along the medio-lateral axis in both OBs for all fish. Values close to 1 indicate lateral locations, whereas values close to 0 indicate medial locations. Note the lateral location of amino-acid selective ipsilateral cells compared to the random distribution indicated by the gray dotted line (***p<0.001; one-sample Student’s t-test). Amino-acid selective cells are similarly located in the contralateral and ipsilateral OBs (ns, non significant; Student’s t-test). (N). Pairwise similarity in mitral cells responses as a function of the distance between them (ipsilateral, n=6 fish; contralateral, n=6 fish; Student’s t-test). Dotted lines indicate average correlation after shuffling the spatial locations of mitral cells. AA: amino acids; pgf2α: prostaglandin 2α; OB: olfactory bulb; ipsi: ipsilateral; contra: contralateral; ns, non significant.

Our anatomical results showed that direct interhemispheric projections between the OBs follow a strict topographical organization and connect similarly tuned glomeruli across hemispheres (Figure 2). Moreover, different odor categories (such as amino-acids, bile acids and pgf2α) were previously shown to specifically activate spatially distinct OB domains [22]. Based on these findings, we hypothesized that interhemispheric connections would communicate odor category specific information across spatially distinct domains between two hemispheres. To test this, we first compared the odor category specificity of the ipsilaterally and contralaterally-evoked mitral cell odor responses. Consistently, we observed odor category selective neurons present both in the ipsilateral as well as in the contralateral OB (Figure 4D,F & H). To quantify the response selectivity of OB neurons, wecalculatedthelifetimesparseness[54] of individual mitral cell odor responses. A neuron with high sparseness value responds primarily to one or a few odors, whereas a neuron with low sparseness value respond with similar amplitude to several odors. We found no difference in lifetime sparseness of mitral cell odor responses in the ipsilateral OB and deafferented contralateral OB (Figure 4I), indicating a similar level of odor category selectivity between ipsi- and contralateral mitral cells. Next, we compared the spatial distribution of odor responses in the ipsilateral and deafferented contralateral OB. Based on visual inspection, odor responses appeared to be mirror-symmetric and thus located in functionally homologous spatial domains (Figure 4J-K). To quantify this mirror-symmetry, we mapped the location of mitral cells selectively responding to distinct odor categories. Since pgf2α activates a restricted set of neurons in the caudo-ventral OB [55], we calculated a caudality index (see methods) measuring the location of pgf2α-selective neurons along the antero-posterior axis. We found that both OBs exhibit similarly high caudality index for pgf2α (Figure 4L). Amino acids activate specifically the lateral zebrafish OB. Hence, we calculated a laterality index to quantify the mirror-symmetry of amino acids selective neurons, and found that both OBs exhibit similarly high laterality index (Figure 4M). To further quantify and compare the spatial distribution of ipsi- and contralaterally evoked odor responses, we calculated the pairwise similarity of mitral cell odor responses, as a function of the distance between them [54]. Both in the ipsi- and contralaterally evoked odor responses, we observed that spatially nearby mitral cells within each OB exhibit similar odor response tuning, which rapidly decreased with increasing distance (Figure 4N).

### Natural odors elicit non-topographically organized responses in the inhibitory interneurons of the contralateral OB

Our dye electroporation of the zebrafish homolog of the olfactory cortex, Dp, showed dense projections that terminated in the granule cell layer of the contralateral OBs (Figure 1E), where inhibitory interneurons (granule cells) are located [56–58]. These results suggest that the two OBs are also indirectly connected through the top-down telencephalic feedbacks, likely recruiting inhibitory interneurons. To test this hypothesis, we labeled the deep layers of the OB with the synthetic calcium indicator Rhod2-AM, and measured the odor responses of the inhibitory interneurons located in the core layers (>50 µm deep, [58]) of the ipsilateral (Figure 5A-B) and deafferented contralateral bulbs (Figure 5C-D). Odor stimulation elicited prominent responses in the inhibitory interneurons on the ipsilateral side (Figure 5E). Moreover, interneurons in the deafferented contralateral OB were also activated, thus confirming our hypothesis (Figure5F).

**Figure 5:**
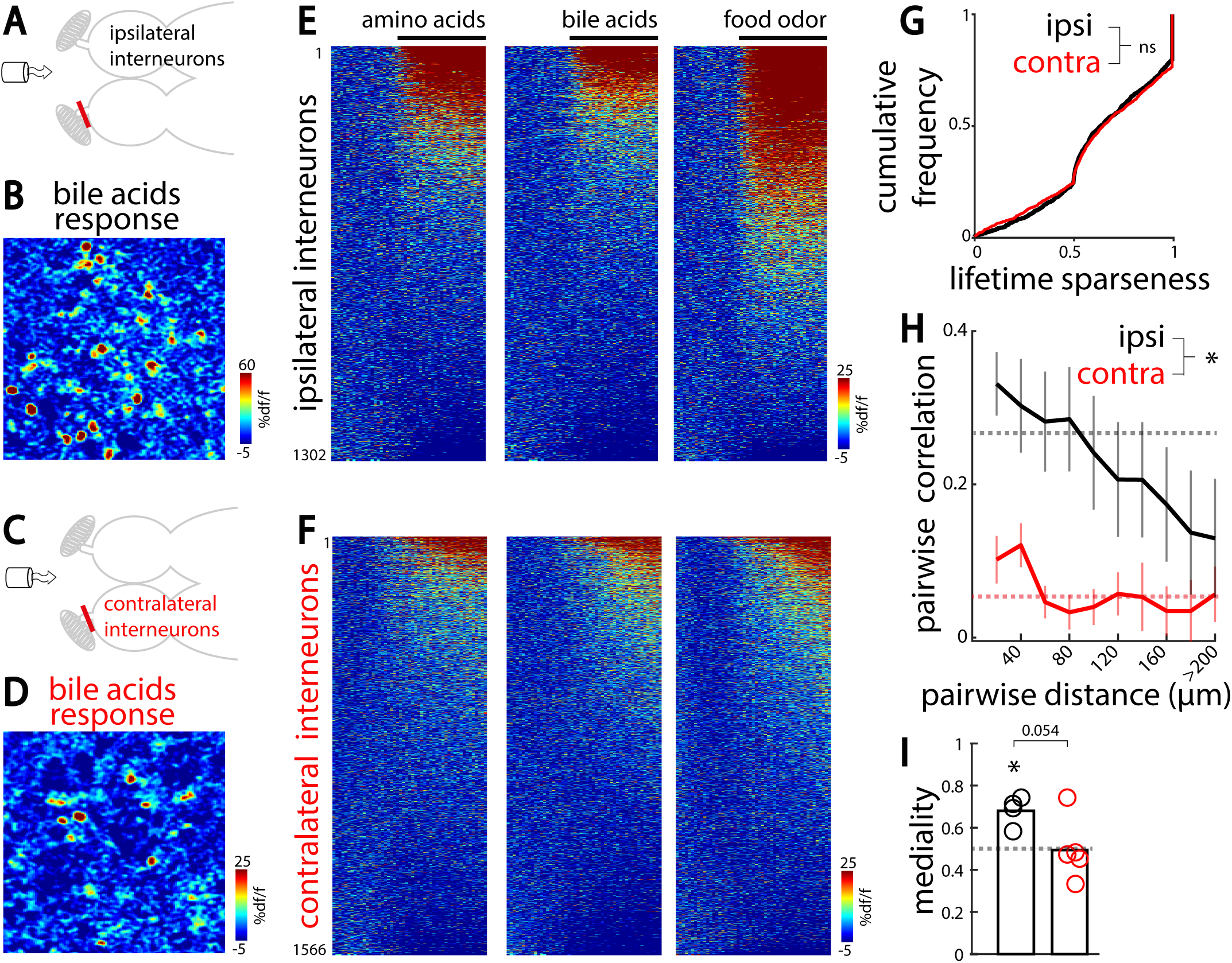
Olfactory bulb interneurons are diffusely activated by the contralateral olfactory inputs. (A). Illustration of the experimental setup depicting an adult zebrafish brain explant, injected with a synthetic calcium indicator (Rhod2 am) labelling inhibitory interneurons. A tube delivers odors to the nostrils. The stimuli used are amino acids, bile acid and food odor extract. The olfactory nerve is sectioned on the contralateral OB. (B). Ipsilateral interneurons response to bile acid. (C). Same setup as in A, except that the deafferented OB contralateral to odor stimulation is imaged. (D). Contralateral interneurons response to bile acid. (E). Odor response time course of all ipsilateral interneurons (1302 cells in 4 fish). For each odor, interneurons are sorted by decreasing responsiveness. (F). Odor response time course of all contralateral interneurons (1566 cells in 5 fish). (G). Cumulative frequency of lifetime sparseness for all contralateral and ipsilateral interneurons (ns, non-significant; two-sample two-tailed Kolmogorov-Smirnov test). (H). Pairwise similarity in interneurons response as a function of the distance separating them (ipsilateral, n=4 fish; contralateral, n=5 fish, *p<0.05; Student’s t-test). Dotted lines indicate average correlation after shuffling the spatial locations of interneurons. (I). Spatial distribution of ipsilateral and contralateral interneurons responding selectively to bile acids, along the medio-lateral axis for all fish. Values close to 1 indicate medial locations, whereas values close to 0 indicate lateral locations. Ipsilateral bile acid-selective interneurons are located medially (* p<0.05; one-sample Student t-test for comparison with random distribution indicated by gray dotted line). Contralateral bile-selective interneurons are located less medially than ipsilateral ones (p=0.054, two-sample two-tailed Student’s t-test with ipsilateral interneurons).

The topographic odor map within the OB is not prominent in the rodent piriform cortex [52,59–61] and in the zebrafish Dp [31,43]: axons from similarly tuned mitral cells project diffusely to these areas without strong spatial preference. Our results add to this by showing that Dp neurons in turn diffusely innervate the contralateral OB granular cell layer (Figure 1). Based on these findings, we hypothesized that the OB interneurons activity pattern recruited by contralateral olfactory inputs would not be topographically organized. To test this, we first compared the odor category specificity of the ipsilaterally and contralaterally-evoked interneuron responses using the lifetime sparseness measure. We observed that odor response tuning of ipsilateral and contralateral interneurons were similar (Figure 5G). Previous studies showed that odors evoke spatially organized responses in zebrafish olfactory bulb interneurons, which partially reflect the topography of mitral cell odor responses [58]. In line with these earlier results, we observed stronger correlations between odor response profiles of nearby interneurons of the ipsilateral OB (Figure 5H). In contrast, odor responses of interneurons in the deafferented contralateral OB exhibited significantly less spatial organization than the ipsilateral interneurons (Figure 5H, contra). Similarly, we observed that symmetric and spatially organized activation of ipsilateral interneurons elicited by bile acids was not present in the contralateral deafferented OB (Figure 5I). These results confirmed our hypothesis and highlight a global and non-topographic recruitment of interneurons through interhemispheric connections, which is mediated through diffuse projections received from the zebrafish homolog of the olfactory cortex, Dp (Figure 1E).

### Interhemispheric connections improve the detection of a reproductive pheromone within a background of olfactory noise

Our results showed that interhemispheric excitation, which originates from the contralateral OB, is spatially organized and recruits mirror-symmetric mitral cells in both OBs. Whereas interhemispheric inhibition, which originates from the innervation of deeper OB layers by the contralateral Dp, is not spatially organized and recruit broadly distributed inhibitory interneurons. Such global inhibition could facilitate interhemispheric suppression and mediate odor source localization or serve as a gain control mechanism. However, strong and glomeruli specific interhemispheric excitation could enhance the sensitivity of the OB circuit, especially for detecting odors that activate few specific glomeruli, without activating the entire network that would otherwise recruit strong global interhemispheric inhibition. Thus, we hypothesized that this balance between strong contralateral focal excitation and global inhibition could improve the detection of odors activating few glomeruli in noisy conditions, where odor detection is challenging, for example in the presence of a background odor simultaneously activating multiple OB glomeruli.

To test this, we compared how mitral cells with or without active interhemispheric connections detect pgf2α, an olfactory cue activating two ventral glomeruli. Pgf2α was either presented alone or together with varying concentrations of food odor, forming odor mixtures (Figure 6A-B). Our functional recordings confirmed that pgf2α activates a restricted set of selectively tuned mitral cells, whereas the competing stimulus food odor activates a large portion of the OB circuits (Figure 6C-F).

**Figure 6:**
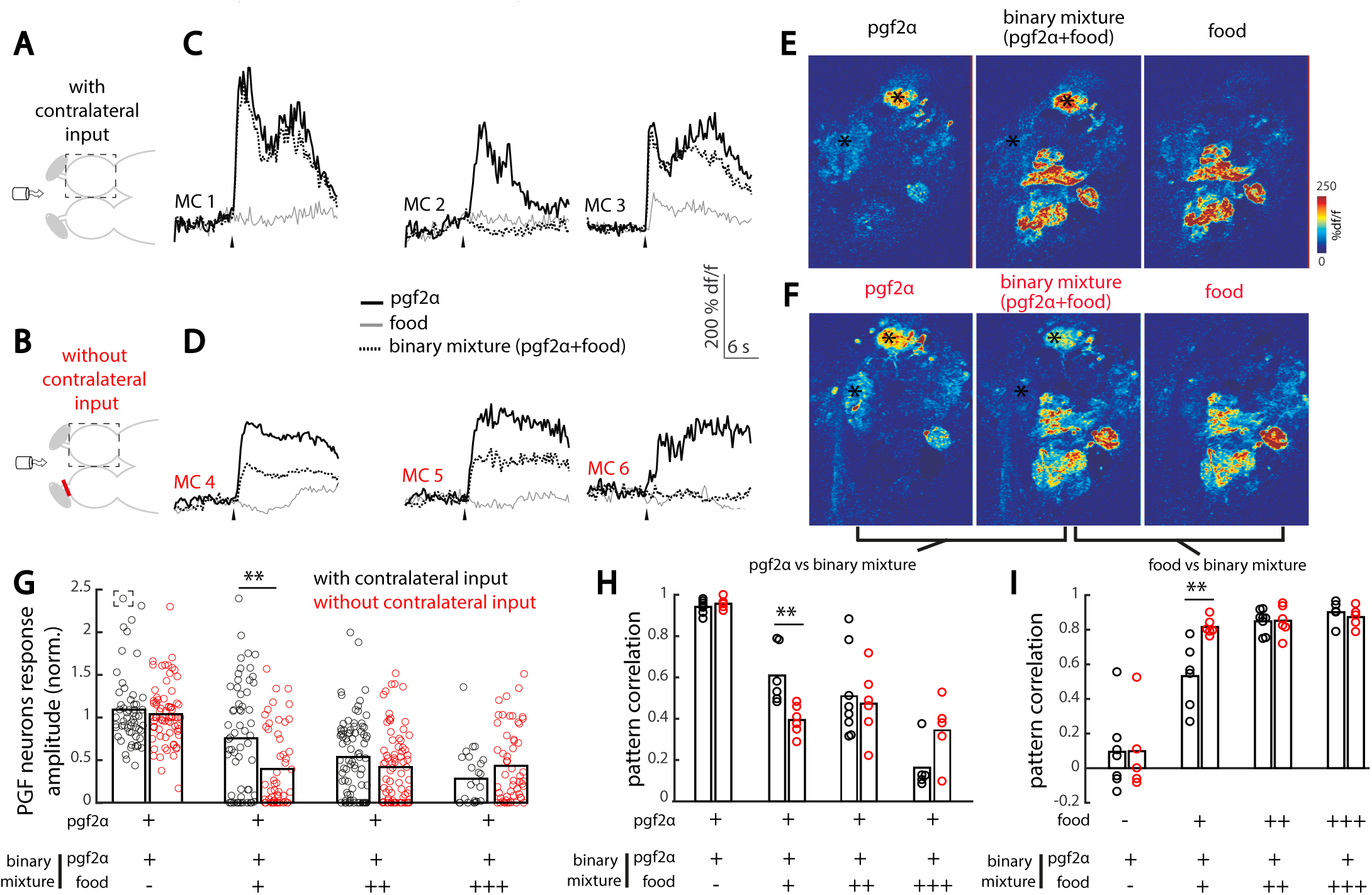
Contralateral olfactory inputs preserve reproductive pheromone detection in food odor background. (A). Illustration of the experimental setup depicting an adult zebrafish brain explant, expressing GCaMP6s in excitatory neurons, and the tube for odor delivery to the intact nose. Three odors were used: pgf2α, food odor extract (food) and a binary mixture of pgf2α and food odor extract. Food odor was delivered at different concentrations (control (-): 0 µg/mL; low (+): 25 µg/mL; medium (++): 100 µg/mL; high (++): 400 µg/mL). The dotted square indicates the imaged olfactory bulb. (B). Ventral fish brain dissection with the contralateral olfactory nerve sectioned, preventing contralateral inputs to modulate the odor responses measured. (C). Odor response time course of three representative pgf2α-selective mitral cells in presence of contralateral olfactory inputs. Black arrowheads indicate odor onset. (D). Odor response time course of three representative pgf2α-selective mitral cells without contralateral olfactory inputs. (E). Odor response maps to pgf2α, food odor and their binary mixture in the olfactory bulb receiving contralateral inputs. Black asterisks indicate pgf2α-selective glomeruli. (F). Odor response maps for the same fish and imaging plane than in E, without contralateral inputs. (G). Relative response of pgf2α-selective cells to the odor mixture in all conditions. Each circle represents a pgf2α-selective mitral cells with (in black) or without (in red) contralateral inputs. The response amplitude of each pgf2α-selective mitral cell to the odor mixture is normalized to its own response to pgf2α alone. Values close and superior to 1 represent conserved or enhanced responses to pgf2α in presence of food odor, respectively. Values close to zero indicate suppression of the pgf2α response in the presence of food odor (**p<0.01, Mann-Whitney U-test). The number of pgf2α-selective mitral cells is as follow with contralateral inputs (control (-): 57 MCs in 7 fish; low (+): 61 MCs in 6 fish; medium (++): 83 MCs in 8 fish; high (+++): 25 MCs in 5 fish) and without contralateral input (control (-): 72 MCs in 5 fish; low (+): 53 MCs in 6 fish; medium (++): 77 MCs in 6 fish; high (+++): 56 MCs in 5 fish). The response of the cell placed in a dashed square is out of range (3.3). (H). Correlation between mitral cells’ odor responses to pgf2α and to odor mixtures for all conditions (**p<0.01, Student’s t-test). High and low pattern correlation values indicate high and low similarities with the pgf2α response pattern, respectively. Each circle represents correlation within a single fish, with contralateral input (in black) or without contralateral input (in red). (I). Correlation between mitral cells’ odor responses to food odor and to odor mixtures for all conditions (**p<0.01, Student’s t-test). High and low pattern correlation values indicate high and low similarities with the food odor response pattern, respectively.

We observed that increasing concentrations of food odor within the binary mixtures gradually suppressed the responses of mitral cells tuned to pgf2α (Figure 6C-D, G). Confirming our hypothesis, we found that this suppression of pgf2α responses was less prominent in the presence of active interhemispheric connections. Indeed, pgf2α-selective mitral cells that received contralateral inputs retained 75% of their initial pgf2α response when pgf2α was presented with low concentration of food odor, versus only 40% without contralateral inputs (Figure 6G). This was likely due to the available interhemispheric excitation that compensated for the global suppression elicited by the food odor.

Next, we asked whether this improved responses of the pgf2α-selective mitral cells to the odor mixture would be sufficient to maintain pgf2α neural response pattern. Increasing the concentration of background food odor presented together with pgf2α gradually reduced the correlations between the pgf2α and mixture neural representations, shifting the mixture activity pattern away from pgf2α (Figure 6H) and towards food odor (Figure 6I). Confirming our hypothesis, we found that the presence of active interhemispheric connections counteracted the degradation of pgf2α signals in the presence of background food odors at low concentrations. Indeed, activity patterns evoked by the odor mixture were more correlated with the pgf2α alone (Figure 6H), and less similar to food odor (Figure 6I) in the presence of active interhemispheric connections. Altogether, our results showed that interhemispheric communication across the OBs can improve the detection of a reproductive pheromone in the presence of background food odor by providing robust odor specific excitation across isofunctional olfactory glomeruli in different brain hemispheres.

## DISCUSSION

In the present study, we found an extensive network of interhemispheric connections between the adult zebrafish OBs with fine topography. We showed that interbulbar projections are present in adult zebrafish OB and directly connect similarly tuned olfactory glomeruli across hemispheres. A link between similarly tuned olfactory glomeruli has previously been described in fruitfly [26], where olfactory receptor neurons send projections to olfactory glomeruli located in both hemispheres. Interestingly, these bilateral projections support rapid orientation towards odor source, through asymmetric neurotransmitter release [26]. In mice, axons of olfactory sensory neurons (OSNs)expressing the same receptor converge onto two glomeruli within the same OB, one located medially and the other laterally [23,62]. Consequently, each bulb in the mouse contains a mirror-symmetric glomerular odor map [23,63] that is absent in zebrafish, in which OSNs expressing the same receptor project onto a single glomerulus [55,64,65]. Interestingly, the murine isofunctional glomeruli within a bulb are specifically connected via an inhibitory intrabulbar circuit [66]. This circuit potentially amplifies the interglomerular activation delay in a concentration-dependent manner, leading the authors to propose that this mammalian circuit linking similarly tuned glomeruli within one bulb enables odorant concentration decoding [67]. These previous findings highlight the interspecies variability of connections linking similarly tuned glomeruli, across and within brain hemispheres. Thus it would be interesting to systematically investigate whether direct connections link the primary olfactory centers (antennal lobes or olfactory bulbs) in different species. Olfactory bulbs neurons are not known to project onto the contralateral OB in the mouse [52,68]. Yet, bilaterally projecting local interneurons (LNs) link both antennal lobes in fruitfly [69], and mitral cells project to the contralateral OB in teleost fish [31,46,70]. A comparative overview of the connectivity between the primary olfactory centers in species adapted to different environmental constraints will certainly provide useful insights into how these interhemispheric olfactory circuits support species-specific behaviors.

The exact role played by interhemispheric OB connections in sensory computation and animal behavior remains unclear. Section of the anterior commissure in mice, which prevents a broad range of interhemispheric interactions including those between olfactory bulbs, resulted in lack of interhemispheric transfer of unilaterally formed olfactory memories [30] or loss of rapid nostril orientation towards odor source [27]. In the present study, we found that interhemispheric olfactory connections elicited a balance of focal excitation and widespread inhibition on the contralateral OB, and thus hypothesized that such a mechanism could improve the detection of focal odors in a background of competing olfactory cues. Indeed, we found that interhemispheric connections rescued the detection of a reproductive pheromone presented together with low concentrations of food odor. This indicates that interhemispheric connections improve the sensitivity of the OB circuit for the reproductive pheromone in challenging detection conditions, such a presence of background olfactory noise, which are likely to be encountered in the fish’s natural habitat. This new finding is particularly relevant since social-related olfactory cues tend to activate a restricted number of glomeruli in the primary olfactory centers of flies and fish [22,55,71], and therefore might be more sensitive to lateral inhibition in presence of competing olfactory cues. Thus, our findings describe a circuit mechanism that serves to enhance the detection of such focal social-related odors when competing background odors are present. It would be interesting to explore in further studies whether this effect also exists for other attractive social-related odors such as bile acids [51,72,73], which also activate a restricted set of ventrally located glomeruli.

In addition to the direct interbulbar projections, we found that the olfactory bulbs are linked via top-down feedback originating from Dp. Top-down feedback from the piriform cortex to the ipsilateral olfactory bulb have been well documented in mammals [46,74], as well as top down feedback from Dp to the ipsilateral OB in fish [74]. In line with this, we observed top-down feedback from Dp to the ipsilateral OB, but we also found that Dp diffusely projects to the contralateral OB granule cells. As could be expected from the loss of chemotopic organization in Dp [31,43] combined with the diffuse nature of their feedback projection to the contralateral OB, we found no obvious topographic organization of contralaterally-evoked inhibition. This is also in line with rodents studies showing that piriform cortex feedback mediates global and non-topographically organized inhibition in the ipsilateral OB [45,75]. It is interesting to note that rodent’s OBs are also linked via a precisely topographically organized multisynaptic circuit: the pars externa of the anterior olfactory nucleus (AONpe) receives input from ipsilateral mitral cells, and in turn projects to the contralateral inhibitory granule cell layer within the isofunctional column [29,30,40]. The contribution of these connections to the contralateral mouse OB activity was recently explored in two studies, one finding evidence of contralateral mitral cell inhibition [30], while another described odor-specific mitral cel excitation and found only weak and randomly distributed inhibition [40]. Thus, our study combined with previous work reveal that a similar interbulbar functional interaction, - i.e. topographically organized focal excitation, potentially combined with broadly distributed inhibition -, could be conserved from fish to rodents and yet mediated by a different circuit architecture.

The direct shared excitation between olfactory hemispheres that we described in this study poses the question of how fish can use inter-nostril delay to rapidly orient towards odor source, as described in sharks [36]. Actually, inter-nostril comparison would work best with segregated and unconnected neuronal pathways that eventually converge to a brain region performing the bilateral comparison. However, the two mechanisms, -i.e. interhemispheric enhancement of odor detection and bilateral comparison of segregated odor inputs -, are not mutually exclusive and could coexist in the same animal. Indeed, others and we have shown that only a subset of mitral cells are linked across hemispheres. In mice, 33% of mitral cells are functionally connected to the contralateral OB [40]. In larval zebrafish, 7 to 56% of the mitral cells project to the contralateral bulb, depending on the glomerulus considered [31]. Our functional results confirm this picture, with around 40% of ipsilateral activity reflected in the contralateral OB (Figure 4G). Thus, the remaining unconnected mitral cells could support inter-nostril comparison for solving odor localization tasks. This could be achieved for example via convergence of these bilateral unconnected cells to a higher brain region integrating inputs from both nostrils, similar to the role played by the AON in rodents [27,39].

Beyond the olfactory system, the present findings add to the existing literature describing the anatomical organization and functional role of interhemispheric connections in sensory processing [9,10]. Evidence of interhemispheric connectivity between early topographic maps in other sensory modalities than olfaction is scarce: a direct interhemispheric connection between retinae, mediated by contralaterally projecting ganglion cells, was recently identified in vertebrates [76]. This connection is only present at perinatal stages and might underlie the synchronization of bilateral retinal waves during development. As interbulbar connections are already present at fish larval stages [31], during which the olfactory maps is established and refined [77], the zebrafish interbulbar pathway might serve a similar role in maintaining the symmetry of bilateral odor maps during development. Yet, the maintenance of this extensive and highly spatially organized circuit throughout adulthood, combined with its role in increased detection of an (adult) sex-pheromone, suggest that this circuit is not just a remnant of embryonic structure, but instead confers a substantial advantage for adult fish in detecting sensory cues important for survival.

Higher cortical sensory areas in vertebrates are linked through interhemispheric fibers, mostly mediating contralateral hemisphere inhibition [78,79]. In this study, we found a dual component inhibitory and excitatory interhemispheric circuit linking homologous sensory maps. More specifically, we found that focal excitatory projections directly activate similarly tuned neurons in the contralateral hemisphere, and indirectly recruit widespread lateral inhibition, eliciting a balance of topographically organized excitation and non-topographic inhibition on the contralateral hemisphere. Taken together with previous studies, our findings indicate that the balance between contralateral excitation and inhibition might vary according to the brain regions connected. A similar variability in contralateral excitation-inhibition balance was also observed between different layers in the mouse somatosensory cortex [80].

Interhemispheric inhibition between cortical regions contributes to hemisphere dominance leading to a lateralization of complex brain functions such as language [81]. In the visual, somatosensory and auditory system, interhemispheric inhibition is also thought to support additional higher order properties such as stimulus location [34,35], and attention[11,12,14]. Here, our functional recordings show focal stimulus selectively enhances the response amplitude of similarly tuned contralateral neurons. Further, we found that the balance of topographically organized excitation and non-topographic inhibition facilitates the detection of a focal stimulus presented in a noisy background. This is reminiscent of interhemispheric sensory computations arising in a specialized region of the auditory cortex of echolocating bats, where focal electrical stimulation of the contralateral cortex selectively enhance the response of similarly tuned neurons specialized in echo delay detection [16]. Thus, we propose that interhemispheric connections might not only play a role in higher cognitive functions such as lateralization, attention and stimulus source localization, but could also modulate sensory gain at early stages of stimulus detection, through a balance of focal excitation and widespread inhibition, and optimize sensory perception.

## ACKNOWLEDGEMENTS

The authors thank Misha Ahrens for sharing Tg(elavl3:GCaMP6s) zebrafish and Yoshihiro Yoshihara for sharing Tg(SAGFF179A:Gal4:UAS:GFP) zebrafish. Discussions with members of the Yaksi lab helped shaping this study and Stephanie Fore, Pradeep Lal and Karoline Hovde provided helpful comments on the manuscript.

## AUTHOR CONTRIBUTIONS

F.K.: Conceptualization, Funding acquisition, Methodology, Data Collection, Data analysis, Writing – original draft, Writing – review & editing E.Y.: Conceptualization, Funding acquisition, Methodology, Resources, Supervision, Writing – review & editing

## ABBREVIATIONS

ACSF: artificial cerebrospinal fluid;
AON: anterior olfactory nucleus;
Dp: dorsal part of the dorso-lateral pallium;
LN: local olfactory interneuron;
lVpG: lateral ventro-posterior glomerulus;
mVpG: medial ventro-posterior glomerulus;
OB: olfactory bulb;
OSN: olfactory sensory neuron;
Pgf2α: prostaglandin 2α.

---

**Supplementary Figure 1.**
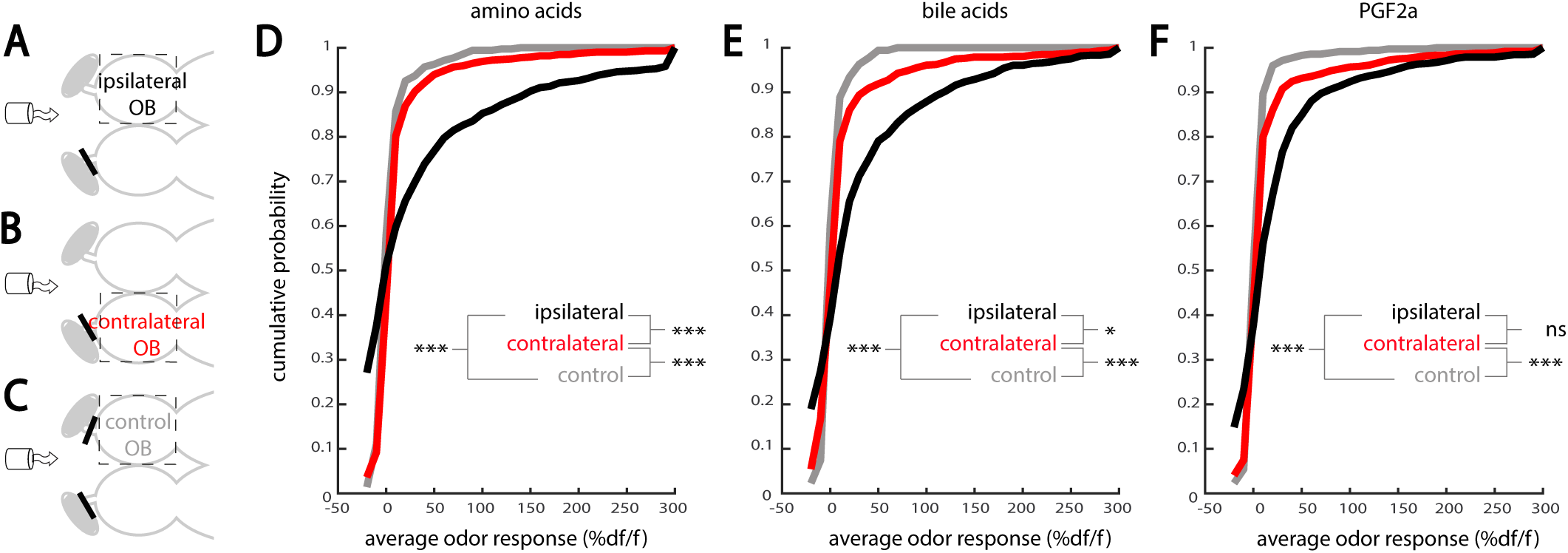
Odor-induced activity on the contralateral OB. (A). Ipsilateral recordings: the olfactory nerve was sectioned on the side contralateral to the imaged OB. (B). Contralateral recordings: same than in A, but responses from mitral cells in the deafferented OB contralateral to odor simulation were recorded. (C). Control recordings: same as in A and B, but with olfactory nerve sectioned bilaterally (control condition). (D, E & F). Cumulative probability distribution of mitral cells response to amino acids, bile acids and pgf2α, respectively (760 ipsilateral cells in 6 animals; 715 contralateral cells in 6 animals; 349 control mitral cells in 4 animals; ***p<0.001, *p<0.05, n.s., non-significant; two-sample two-tailed Kolmogorov-Smirnov test). OB: olfactory bulb; pgf2a: prostaglandin 2α.

## METHODS

### Experimental animal model

Experiments were conducted in adult zebrafish (Danio rerio) of both sexes, aged 8-14 months. The following transgenic lines were used: Tg(SAGFF179A:Gal4:UAS:GFP) [51]; Tg(elavl3:GCaMP6s) [53] and nacre/mitfa. Animals were kept under standard laboratory conditions (28.5°C; 14/10 hour light/dark cycle). All procedures involving animals were approved and followed the animal care guidelines issued by the government of Belgium and the Norwegian Food Safety Authorities (FOTS 19910).

### Preparations of brain explants

Nose-attached brain explant. The experiments were conducted in an ex vivo nose-attached brain explant preparation [22,42,43]. Adult zebrafish were euthanized by immersion in ice-cold water, then decapitated to ensure death. The head was transferred in cold artificial cerebrospinal fluid (ACSF) bubbled with carbogen (95% O2/5% CO2). The ACSF was composed of the following chemicals diluted in reverse-osmosis purified water: 131 mM NaCl, 2 mM KCl, 1.25 mM KH2PO4, 2 mM MgSO4.7H2O, 10 mM glucose, 2.5 mM CaCl2 and 20 mM NaHCO3. The eyes, jaws and ventral part of the skull were carefully removed using forceps, exposing the ventral OBs and telencephalon, and leaving the nose attached. The brain explant was then affixed using tungsten pins to a small petri dish coated with Sylgard (World Precision Instruments) and left to equilibrate at room temperature for >30 min, under constant ACSF perfusion. Depending on the experiment, the olfactory nerve was unilaterally, or bilaterally, sectioned between the olfactory epithelium and the rostral end of the OB, using a sharpened insect pin. Complete deafferentation was verified under a binocular microscope.

Bolus injection of calcium indicator. To record activity in interneurons, a synthetic calcium indicator was injected in the OB [42]. Rhod-2-AM (50 µg, Invitrogen, CAT # R1245MP) was suspended in 16 µL of dimethyl sulfoxide-pluronic (Invitrogen, CAT: P3000MP) and kept at 4°C. Aliquots of the stock solution were diluted (1/10) in ACSF the day of the experiment and gently pressure-injected using a pulled glass micropipette. Experiments started >30 min after the last injection to allow for dye uptake in neurons and clearance from intercellular space.

### Olfactory stimuli

We used electrical stimulation of the olfactory nerve, and bath application of odorants to the intact nose, to elicit reproducible olfactory responses. For experiments with unilateral stimulation with odorants and microstimulation on the contralateral side, a small custom-made polystyrene separator was inserted in front and below the nose to prevent the odorant to reach the contralateral nostril (Figure 3).

Olfactory nerve microstimulation. Microstimulation was performed using pulled glass micropipettes (∼10µm tip diameter) filled with ACSF and containing a silver wire connected to a computer-controlled stimulus isolator (Isoflex amplifier, AMPI). The tip of the micropipette was inserted in the olfactory nerve and a train of short current pulses were applied during 3 seconds (pulse duration: 50 ms; 5-10-20 μA for low, medium and high intensity simulation, respectively).

Odorants. All odorants were purchased from Sigma Aldrich. Odor stocks were prepared in reverse osmosis purified water and kept as aliquots at −20°C. Aliquots of stock solutions were diluted to their final concentration in ACSF on the day of the experiment. Amino acids mix contained arginine, asparagine, aspartic acid, alanine, phenylalanine, histidine and methionine, each diluted at 10-4 mol/L. Bile acid mix contained taurodeoxycholic and taurocholic acids, each diluted at 10-4 mol/L. Prostaglandin 2⊠ was used at the final concentration of 5 10-7 mol/L, alone or within binary mixture with food odor. Food odor stock was prepared using commercially available fish food (SDS100, Scientific Fish Food): 1 g of food was incubated in 100 mL of ACSF and filtered. Food odor stock was diluted 1/50 in ACSF for all experiments, except for the one displayed in Figure 6, where low (dilution 1/400), medium (dilution 1/100) and high (dilution 4/100) concentrations were used. A computer-controlled low pressure injection valve (Valco Instruments, C22Z-3186EUH) introduced the odorants in the perfusion stream (3 mL/min) delivered through a tubing positioned 1 mm in front of the nostrils. The tubing was made of non-adherent teflon (Teknolab, 0.8mm internal diameter) and was rinsed with reverse osmosis purified water after each trial to avoid contamination between successive stimuli. Multiple odorants were delivered in a pseudo-randomized order and separated by at least 2 min in order to avoid neural habituation.

### Axonal projection tracing using dextran-coupled dyes

For axonal projection tracing, neurons were loaded with an fluorescent tracer using local electroporation as described in [52]. The dura covering the brain explant was carefully removed at the labeling sites using flattened-tip forceps to facilitate the dye electroporation. A solution of tetramethylrhodamine-dextran (TMR-dextran, 3000 kDa, Invitrogen, CAT: D3306) diluted at 12 mg/mL in phosphate buffer saline (VWR, CAT: 97062-730) was aliquoted and stored at −20°C to be thawed just before the experiment. Pulled-glass micropipettes were backfilled with 2 µL of dye and the remaining space was filled with ACSF. A silver wire was inserted in the micropipette and connected to a stimulus isolator (ISO-Flex, A.M.P.I.). The micropipette tip was directed to the center of the area of interest under a two-photon microscope and trains of current pulses (500 pulses of 30-40 µA, 25 ms duration, 2Hz) were delivered. The brain explant was bathed at room temperature in a flow of ACSF bubbled with carbogen for at least 6h to allow the dye migration along the neurites. We used a two-photon imaging system (LSM 7 MP upright with 20× water immersion objective, Zeiss) to take anatomical scans of the brain, since it enabled access to deep structures and reduced fluorophore bleaching. A mode-locked Ti:Sapphire laser (MaiTai Spectra-Physics) tuned to 840 nm was used for combined excitation of GFP and TMR. Labeled contralateral neurites were then reconstructed by an experimenter blind to the location of the ipsilateral electroporation sites based on the TMR labeling (red channel) in Neuron Studio [82] or using the Simple Neurite Tracer plugin [83] in ImageJ. Neurites reconstructed were then superimposed on the stack acquired in the GFP channel that provided the OB outline.

### Two-photon calcium imaging

Neural activity measurements were collected with two-photon laser scanning systems. For the experiment described in Figure3, data were collected for 1-3 planes per fish, using successive recordings of a single plane at 3.4Hz (LSM7MP upright with 20× water immersion objective, Zeiss). For the remaining experiments, volumetric images were recorded at 3.3Hz (upright with 16× water immersion objective, Scientifica) for 6 planes simultaneously, which were evenly spanning the ventral side of the OB within a range of 10 to 110 µm deep. In both cases, a mode-locked Ti:Sapphire laser tuned to 920 nm (for GCaMP6s recordings) or 840 nm (for Rhod2AM recordings) was used to excite the fluorophores.

### Calcium imaging analysis

Recordings were corrected for movement in Matlab using a modified version of the algorithm used in [84,85]. Recordings were then visually inspected and discarded whenever tissue drift was observed. Individual neurons were manually segmented and the corresponding raw fluorescent traces were calculated by averaging the value of all pixels belonging to a given neuron for each time point. The change in fluorescence (% df/f) relative to pre-stimulus baseline was calculated as follow:

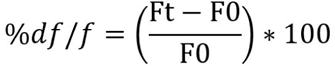

where F0 is the averaged 6 seconds of pre-stimulus fluorescence for each neuron and Ft the fluorescence of a neuron at time t. The stimulus response of each neuron was calculated by averaging the %df/f during the 10 seconds post-stimulus onset (for odor responses) or 3 seconds post-microstimulation onset (for contralateral olfactory nerve microstimulations). Images of stimulus responses within an entire recording plane were obtained by applying the same procedure to every pixel (instead of neuron) and smoothening the resulting image using a Gaussian filter. To calculate the selectivity of odor response of the entire neuronal population, lifetime sparseness was calculated as follow [86,87]:

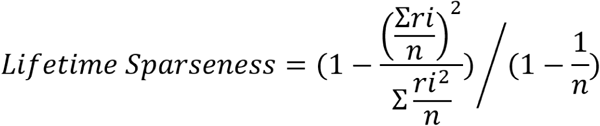

where ri is the response of an individual neuron to the ith odor and n is the total number of odors. Cells with negative responses to all odors were not included in the calculation (remaining mitral cells: 628 ipsilateral cells in 6 fish; 606 contralateral cells in 6 fish; remaining interneurons: 1196 ipsilateral interneurons in 5 fish; 1466 contralateral interneurons in 4 fish). Low values indicate low selectivity whereas values close to 1 indicate that the neuron respond to only a small fraction of the odor set tested. The thresholds for detecting a response were set to 70% df/f for mitral cells and 10% df/f for interneurons. To quantify chemotopy, the spatial location of neurons responding to a single stimulus (odor-selective neurons) was allocated to anterior versus posterior OB halves or to medial versus lateral OB halves. Lateral, caudal or mediality indices were then calculated in each animal as the ratio of neurons located in the lateral, posterior or medial OB halves, respectively.

### Statistical analysis

Data was examined for normality using a Shapiro-Wilk test. The significance threshold was set to p value < 0.05. Data were then analyzed using parametric (Student’s t-test) or non-parametric tests (Wilcoxon signed-rank). Two-sample two-sided Kolmogorov-Smirnov test were used for population distribution. Data is represented as mean +/- standard error of the mean unless stated otherwise.

